# *Klebsiella taiwanensis* sp. nov., and *Pseudomonas ficus sp. nov.*, novel taxa isolated from figs (*Ficus*)

**DOI:** 10.64898/2026.11.23.768557

**Authors:** Samuel Miller, Austin C. Link, Nam Lu, Kimberly A. Moser, John Wang, Paul A. Lawson, Gavin C. Woodruff

## Abstract

Two bacterial strains, WOUb02^T^ and WOUb67^T^, were isolated from fresh figs (*Ficus* spp.) collected from the island of Taiwan. Cells of WOUb02^T^ and WOUb67^T^ were Gram-stain-negative rods. Strain WOUb02^T^ was non-motile, whereas WOUb67^T^ exhibited motility. Physiologically, both strains grew optimally at 0.5 % (w/v) NaCl and pH 8.0; WOUb02^T^ grew optimally at 37°C, whereas WOUb67^T^ grew optimally at 30°C. Based on 16S rRNA gene sequence similarity, the closest relatives to strains WOUb02^T^ and WOUb67^T^ were *Klebsiella terrigena* ATCC 33257^T^ (98.2%) and *Pseudomonas putida* NBRC 14164^T^ (99.9%) respectively. Subsequent phylogenetic analysis based on 16S rRNA gene sequences revealed that strains WOUb02^T^ and WOUb67^T^ formed distinct lineages within the genera *Klebsiella* and *Pseudomonas*, respectively. Further genomic analyses revealed that WOUb02^T^ and WOUb67^T^ shared average nucleotide identity and digital DNA-DNA hybridization values <95.0% and <70.0%, below the currently accepted thresholds for species-level delineation. Based on phenotypic and genotypic data, strains WOUb02^T^ and WOUb67^T^ could be distinguished from other species with validly published names and are considered to represent novel species in the genera *Klebsiella* and *Pseudomonas*, respectively, for which the names *Klebsiella taiwanensis* sp. nov., and *Pseudomonas ficus* sp. nov., are proposed. The type strains are WOUb02^T^ (=NRRL-B 65719^T^ =CCUG 77855^T^) and WOUb67^T^ (=NRRL-B 65720^T^ =CCUG 77856^T^).

## Introduction

The genus *Klebsiella* within the family *Enterobacteriaceae* has historically encompassed Gram-stain-negative, non-motile, facultatively anaerobic bacteria [1]. According to the List of Prokaryotic Names with Standing in Nomenclature (LPSN) (https://doi.org/10.83108/rn.515890, August 2026), there are 19 *Klebsiella* spp. with validly published names. *Klebsiella* spp. have long been associated with human pathogenesis [2, 3], particularly members of the *Klebsiella pneumoniae* complex [4, 5]. *Klebsiella* are often recovered from clinical samples [6–8], environmental samples [9, 10], and even plants [11, 12].

The genus *Pseudomonas* belongs to the family *Pseudomonadaceae* and consists of Gram-stain-negative, predominantly motile, aerobic species [13]. There are 384 *Pseudomonas* spp. with validly published names according to LSPN (https://doi.org/10.83108/rn.517405, August 2026). While significant clinical interest exists in *Pseudomonas aeruginosa* [14, 15] due to its pathogenicity, *Pseudomonas is* a highly diverse and ubiquitously distributed genus whose members exhibit an exceptionally high level of metabolic diversity [16].

In a continuing study of fig-nematode interactions [17, 18], bacteria from fresh figs (*Ficus* spp.) were isolated and fed to *Caenorhabditis inopinata,* the closest known relative of *Caenorhabditis elegans*. *C. elegans* is a bacterivore that thrives on rotting plant detritus [19–22] and demonstrates increased fitness when reared on bacteria from its natural environment [21, 23]. *Caenorhabditis inopinata*, reproduces in the fresh figs of *Ficus septica* [24, 25]. Using a polyphasic taxonomic approach, two novel bacterial strains, WOUb02^T^ and WOUb67^T^, belonging to the genera *Klebsiella* and *Pseudomonas,* were recovered. The names *Klebsiella taiwanensis* sp. nov., and *Pseudomonas ficus* sp. nov., are herein proposed.

### Isolation and Ecology

Between August 2019 and August 2023, fresh figs (*Ficus*) samples from Taipei, Taiwan (25.0314672, 121.5749889) were collected and subsequently dissected in a sterile Petri dish containing 4 ml of sterile M9 buffer [26]. Following dissection, 200μl of the fig/buffer suspension was spread-plated onto sterile Luria-Bertani (LB) agar and nutrient agar (NA) plates and left at room temperature. These plates were transferred to the University of Oregon, where monocultures were established by multiple plate streaks until pure isolates were obtained. In total, 44 strains were isolated and screened using near-full-length 16S rRNA gene sequencing [17]. Two resulting isolates, WOUb02^T^ and WOUb67^T^, were transferred to the University of Oklahoma for further investigations. After primary isolation, isolates were maintained under aerobic conditions on LB, NA, or tryptic soy agar (TSA) at 25-37°C and preserved in LB broth with 20% (v/v) glycerol at −80 °C.

### 16S rRNA Gene Phylogeny

Strains WOUb02^T^ and WOUb67^T^ were sent to GENEWIZ (South Plainfield, NJ, USA), and near full-length 16S rRNA gene sequences were obtained using Sanger sequencing. Using the EzBioCloud server (www.ezbiocloud.net), the 16S rRNA gene sequences of WOUb02^T^ and WOUb67^T^ were compared to those of the most closely related type strains, as determined by a BLAST search against the GenBank database using EzBioCloud’s Identify feature. The 16S rRNA gene sequences of WOUb02^T^, WOUb67^T^, and closely related species were identified with the EzBioCloud server [27], downloaded, and aligned using Muscle [28]. Phylogenetic trees using the maximum-likelihood method [29] and the Tamura-Nei substitution model [30] were reconstructed with 1,000 bootstrap replications. The phylogenetic trees were constructed using MEGA 12 [31].

The results of 16S rRNA gene phylogenetic analyses indicated strains WOUb02^T^ and WOUb67^T^ were most closely associated with *Klebsiella terrigena* ATCC 33257^T^ (98.2%) and *Pseudomonas putida* NBRC 14164^T^ (99.9%), respectively. The 16S rRNA gene sequence similarity between WOUb02^T^ and its close relatives was below the currently accepted threshold of 98.6 %, which is now routinely used to demarcate this taxonomic rank [32]. Additionally, the topologies of the associated phylogenetic trees constructed using the maximum-likelihood method [29] demonstrate that WOUb02^T^ and WOUb67^T^ (Figs. 1 and 2) cluster within the genera *Klebsiella* and *Pseudomonas*, respectively. Further analyses were conducted for both strains to establish a more robust species-level taxonomic differentiation, especially for WOUb67^T^, as 16S rRNA gene sequences have been shown to convey low discriminatory power for *Pseudomonas* spp. [33].

**Figure 1.**
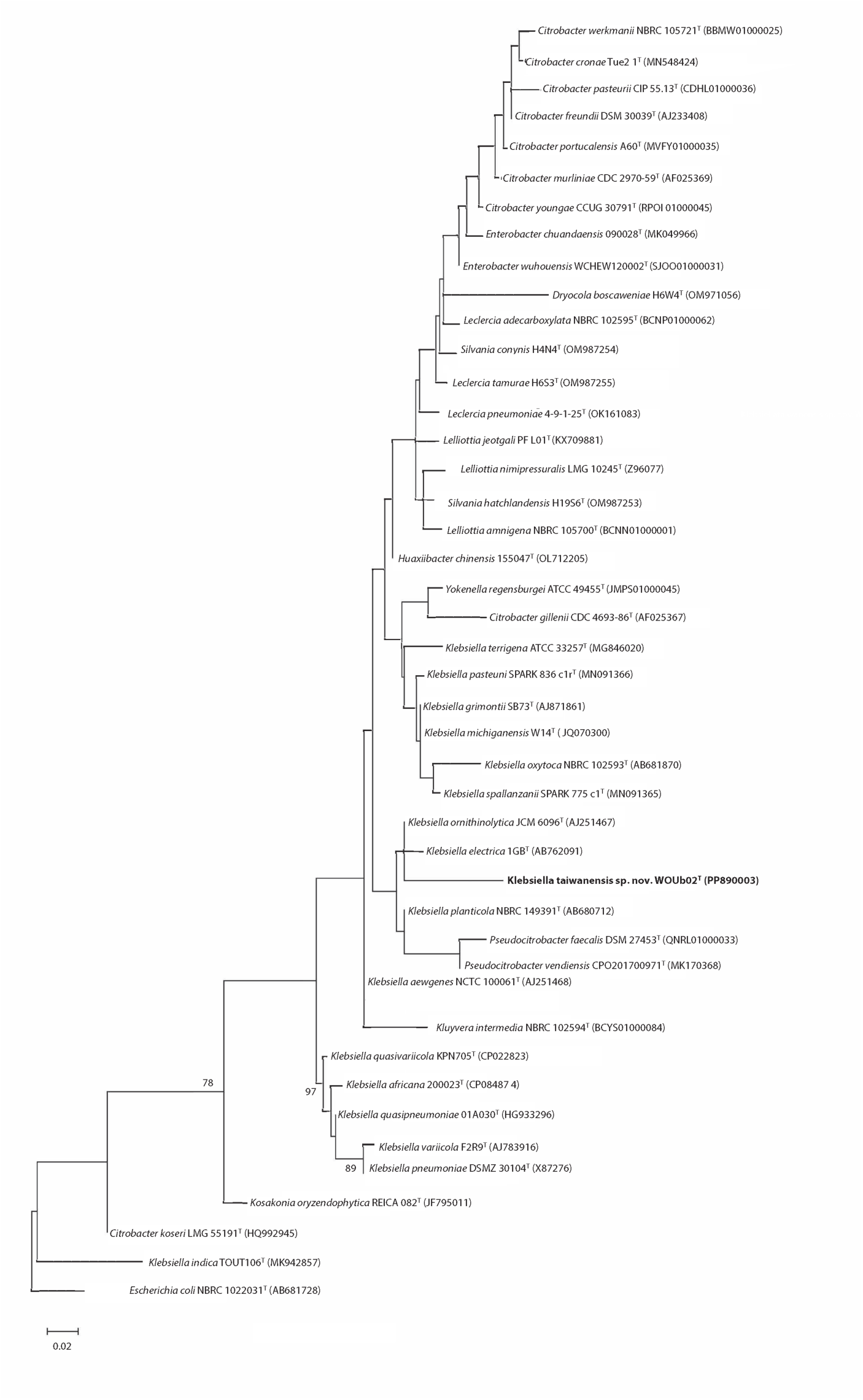
Phylogenetic tree based on 16S rRNA gene sequences showing the relationship between WOUb02^T^ and other members of the genus *Klebsiella*. *Escherichia coli* NBRC 102203^T^ (AB681728) was used as an outgroup. Numbers at the nodes represent bootstrap percentages from 1,000 replicates; only values >70% are shown. The bar indicates nucleotide substitutions per site, 0.02.

**Figure 2.**
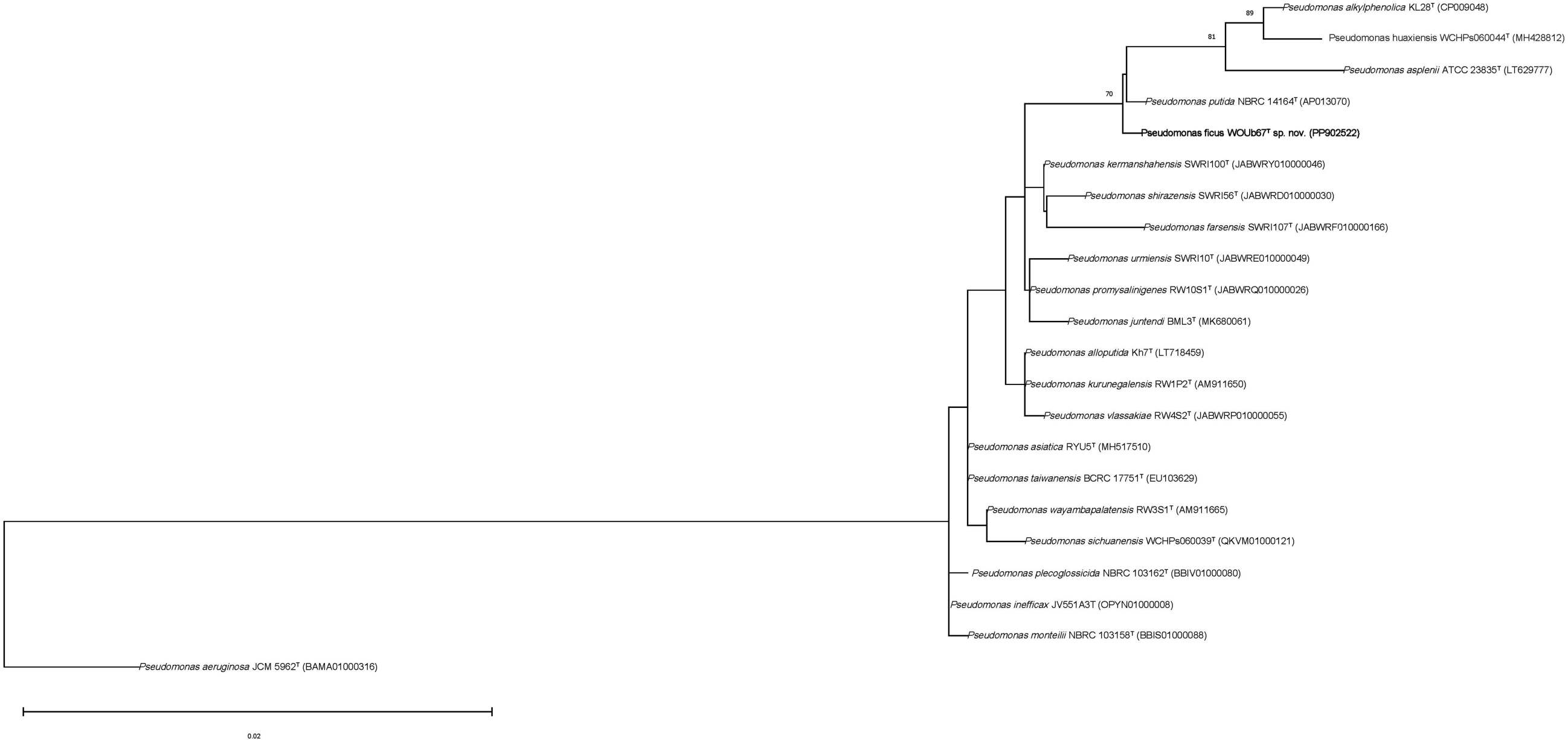
Phylogenetic tree based on 16S rRNA gene sequences showing the relationship between WOUb67^T^ and other members of the *Pseudomonas*. *Pseudomonas aeruginosa* JCM 5962^T^ (BAMA01000316) was used as an outgroup. Numbers at the nodes represent bootstrap percentages from 1,000 replicates; only values >70% are shown. The bar indicates nucleotide substitutions per site, 0.02.

### Whole genome sequencing and phylogenomic analyses

To further clarify the taxonomic relationship between WOUb02^T^, WOUb67^T^, and closely related taxa, the genomes of WOUb02^T^ and WOUb67^T^ were subjected to whole-genome sequencing. Cell biomass for all strains was harvested from overnight cultures and stored in DNA/RNA Shield (Zymo, Irvine, CA, USA). DNA extraction, sequencing, and annotation were performed by a commercial provider (Plasmidsaurus, Eugene, OR, USA) as previously described [34]. Assembly completeness and contamination were assessed using CheckM v1.2.2 [35]. Genome annotation was performed using the NCBI Prokaryotic Genome Annotation Pipeline v6.7 [36].

To delineate species boundaries between WOUb02^T^, WOUb67^T^, and closely related taxa, overall genomic relatedness indices (OGRIs) were calculated so that the most recent proposed minimal standards for the use of genome data for the taxonomy of prokaryotes [30] were used. As such, average nucleotide identity (ANI) [27, 38] and digital DNA-DNA hybridization values (dDDH) [32, 33] were computed. Values for ANI were calculated using EzBioCloud’s ANI Calculator, which uses the OrthoANIu algorithm [27, 38]. The Genome-to-Genome Distance Calculator [39, 40] v3.0 was used to determine dDDH. Additional whole-genome-based phylogenetic analysis of WOUb02^T^ and WOUb67^T^ was performed using the Codon Tree Method [11] (Figs. 3 and 4) within a custom workspace on the Bacterial and Viral Bioinformatics Resource Center (BV-BRC). Visualization was accomplished with the Interactive Tree of Life v6 [41].

**Figure 3.**
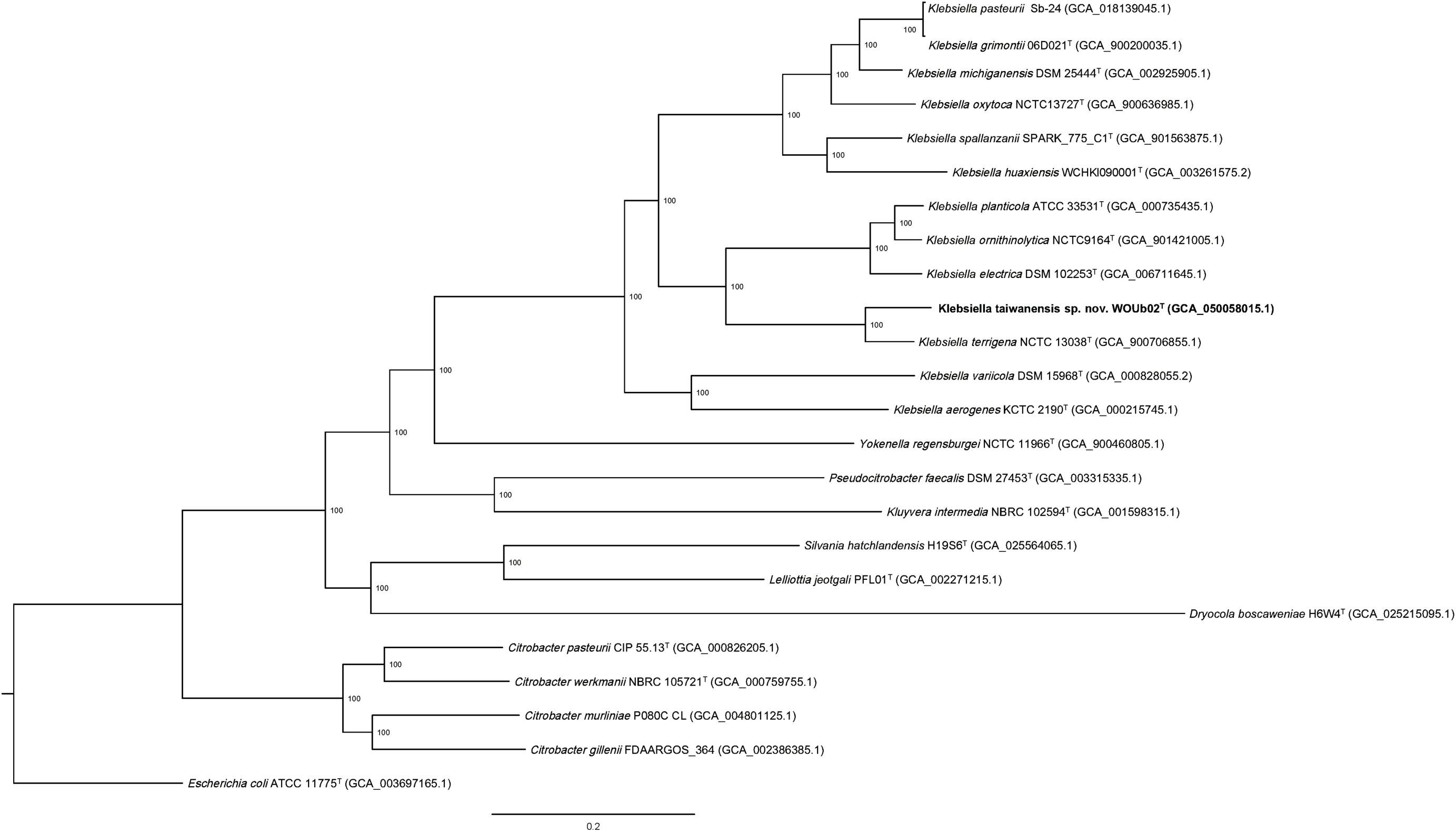
Core-genome phylogenomic tree using a concatenated alignment of 1,000 single-copy genes showing the relationship among WOUb02^T^ among other species of the genus *Klebsiella*. *Escherichia coli* ATCC 11775^T^ (GCA_003697165.1) was used as an outgroup. Numbers at the nodes represent bootstrap percentages from 100 replicates; only values >70% are shown. The bar indicates the mean number of substitutions per site, 0.2.

**Figure 4.**
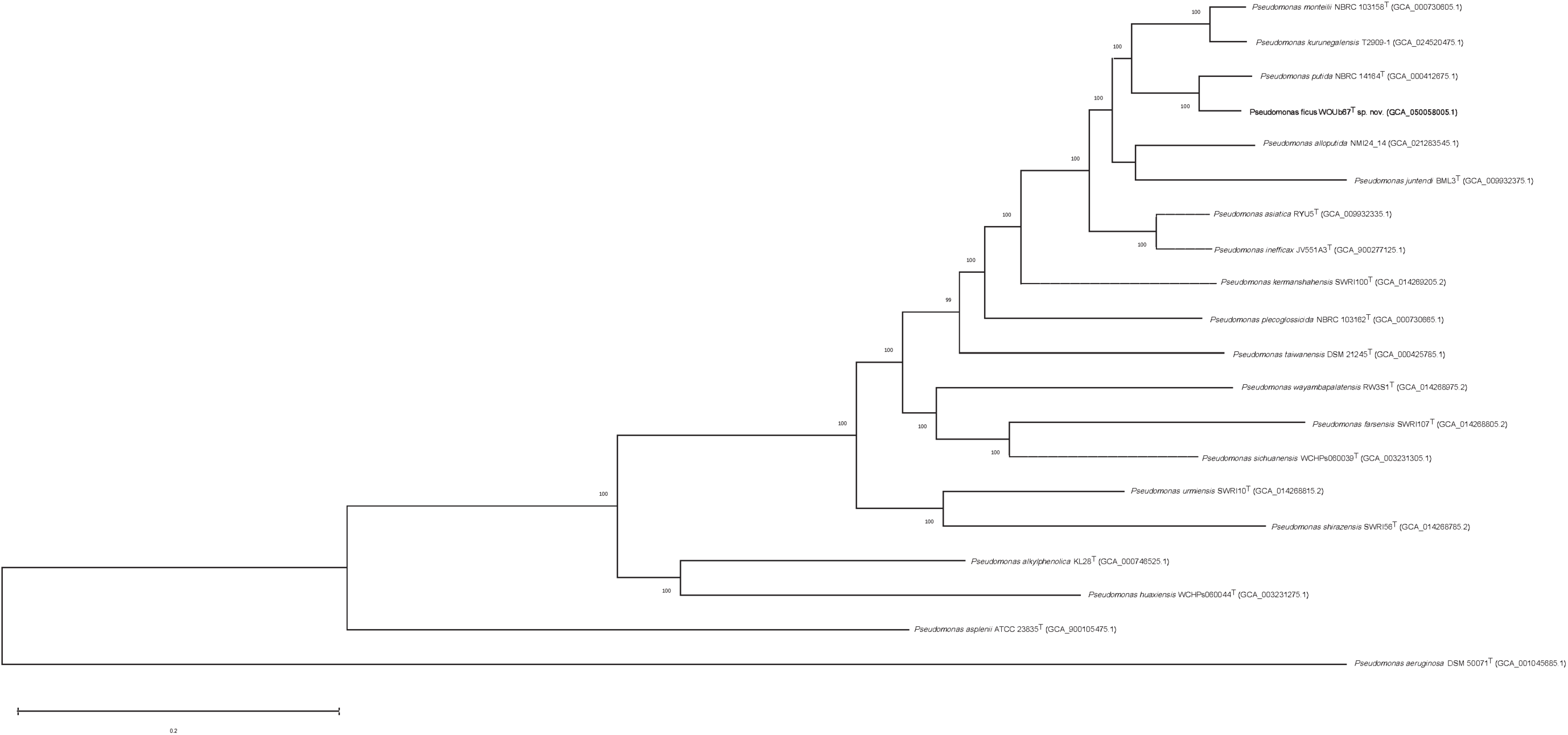
Core-genome phylogenomic tree using a concatenated alignment of 1,000 single-copy genes showing the relationship among WOUb67^T^ and other species of the genus *Pseudomonas*. *Pseudomonas aeruginosa* DSM 50071^T^ (GCA_001045685.1) was used as an outgroup. Numbers at the nodes represent bootstrap percentages from 100 replicates; only values >70% are shown. The bar indicates the mean number of substitutions per site, 0.2.

Once assembled, the draft genome sequence of WOUb02^T^ and WOUb67^T^ consisted of 9 and 2 contigs, respectively. Genome lengths (Mbps) of 6.0 and 6.6 were observed for WOUb02^T^ and WOUb67^T^, respectively. The G+C content (%) was 57.0% and 62.5%, respectively. Genome quality metrics from the NCBI PGAP CheckM analysis (v1.2.2) indicated completeness of 93.6% and 100.0%, whereas contamination levels of 2.7% and 0.2% were observed for WOUb02^T^ and WOUb67^T^, respectively. As such, genomic sequences were deemed acceptable for inclusion in taxonomic and phylogenomic studies [35]. Of the 6,037 predicted genes for WOUb02^T^, 4,870 were protein-coding sequences (CDSs), 25 rRNA genes, and 88 tRNAs were identified. For WOUb67^T^, 5,826 genes were predicted, of which 5,667 were CDSs, 22 rRNA genes, and 76 tRNAs were identified.

The phylogenomic relationships between WOUb02^T^ and its relatives (Fig. 3) complemented the 16S analysis, revealing that WOUb02^T^ was nested within the genus *Klebsiella*, clustering most closely with *K. terrigena* NCTC 13038^T^ (GCA_900706855.1). Similarly, phylogenomic analysis (Fig. 5) showed that WOUb67^T^ is most closely related to *P. putida* NBRC 14164^T^ (GCA_000412675.1) and is nested within the genus *Pseudomonas*. Despite the high 16S rRNA gene similarity value with *P. putida*, ANI and dDDH values of 94.3% and 58.7%, respectively are below the proposed species-level delineation thresholds [39, 42] (<95.0% and <70.0%, respectively) and clearly demonstrate that WOUb67^T^ represents a novel species within the genus *Pseudomonas*.

Therefore, phylogenomic analysis confirmed the affiliation of strains WOUb02^T^ and WOUb67^T^ with the genera *Klebsiella* and *Pseudomonas*, as well as their distinction from all other described species within these two genera (Figs. 3 and 4), consistent with the analyses of phylogenies constructed using 16S rRNA gene sequences (Figs. 1 and 2). The OGRIs for species delineation revealed ANI and dDDH values below the proposed species-level delineation thresholds [39, 42] (<95.0% and <70.0%, respectively). The main genotypic differences between WOUb02^T^, WOUb67^T^, and their close relatives are presented in Tables 1 and 2. Collectively, the comparative OGRIs and phylogenomic analyses strongly indicate that both strains represent two novel, distinct species within the genera *Klebsiella* and *Pseudomonas*.

**Table 1.** Genotypic differential characteristics of WOUb02^T^ and close relatives.

| **Characteristic** | **1** | **2** | **3** | **4** | **5** |
|---|---|---|---|---|---|
| **16S rRNA gene accession** | PP890003.1 | Y17658 | AJ251467 | JMPP01000074 | AB762091 |
| **16S rRNA gene similarity (%)** | 100.0 | 98.3 | 98.3 | 98.2 | 98.2 |
| **Genome accession** | GCA_055451655.1 | GCA_006539725.1 | GCA_901421005.1 | GCA_000735435.1 | GCA_006711645.1 |
| **OrthoANIu (%)** | 100.0 | 94.0 | 85.7 | 85.6 | 85.7 |
| **dDDH (%)** | 100.0 | 55.0 | 30.2 | 30.1 | 30.5 |
| **DNA G+C (%)** | 57.0 | 57.5 | 55.5 | 55.8 | 55.1 |
| **Genome Size (Mbps)** | 6.0 | 5.5 | 5.6 | 5.7 | 5.8 |
| **Number of Contigs** | 9 | 55 | 2 | 83 | 6 |
| **Number of CDSs** | 4,870 | 5,037 | 5,043 | 5,189 | 5,308 |

Strains: 1, WOUb02<sup>T</sup>; 2, *K. terrigena* NBRC 14941<sup>T</sup>; 3, *Klebsiella ornithinolytica* JCM 6096<sup>T</sup>; 4, *Klebsiella planticola* ATCC 33531<sup>T</sup>; 5, *Klebsiella electrica* 1GB<sup>T</sup>.

**Table 2.**
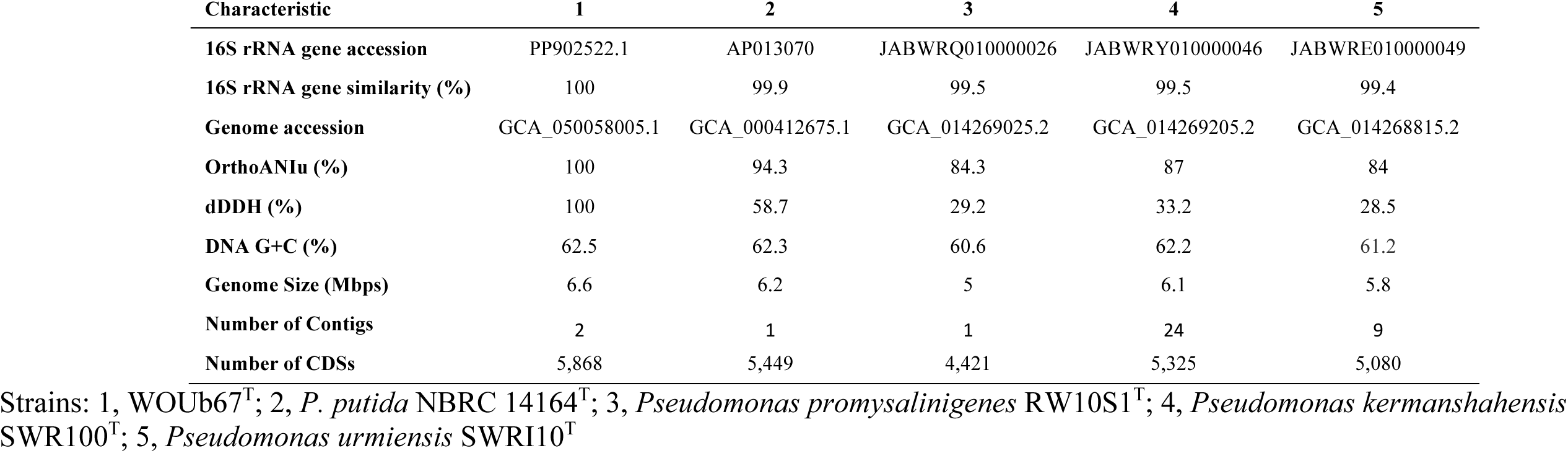
Genotypic differential characteristics of WOUb67^T^ and close relatives.

### Functional genome and *in-silico* analyses

Carbohydrate-active enzyme (CAZyme) analysis was performed using the dbCAN3 server [43, 44]. The genomic repertoire of WOUb02^T^ reflected a high degree of fitness for both the ripening and decomposing stages of figs. The presence of degradation pathways for fructose, mannose, and starch aligns with the carbohydrate-rich environment of ripening figs. Furthermore, its ability to degrade aromatic secondary metabolites via benzoate and homoprotocatechuate pathways suggests a role in breaking down complex plant organic matter. Strain WOUb02^T^ exhibits a CAZyme profile (293 CAZymes) heavily enriched in glycoside hydrolases targeting starch (GH13), fructose (GH32), and pectin (GH28/CE8). The genomic repertoire of WOUb67^T^ similarly supports high fitness within the fig ecology, though with a distinct emphasis on environmental persistence and the degradation of structural polysaccharides. The increased presence of starch-utilization enzymes, including 29 copies of GH13 alongside CBM41 and CBM48 modules, suggests an enhanced capacity to harvest α-glycans. While it can degrade complex aromatics via benzoate and homoprotocatechuate pathways, WOUb67^T^ has a CAZyme profile (296 CAZymes) further characterized by GH94 (cellobiose phosphorylase) and families targeting bacterial and plant-derived glycans (GH23, GH24). Ultimately, the metabolic specialization of WOUb02^T^ and WOUb67^T^ likely facilitates the rapid colonization of ripening fruit and the conversion of botanical sugars into a nutritionally dense biomass. These functional traits may affect the fitness of other fig community members, including bacterivorous nematodes such as *C. inopinata*.

### Ecological Distribution Patterns

Ecological distribution and habitat preference analysis based on the 16S rRNA gene sequence data was computed using the Protologger server v 22.05 (http://www.protologger.de) [45] was used to determine the prevalence and average relative abundance of WOUb02^T^ and WOUb67^T^ across 19,000 amplicon sequencing datasets [46].

The 16S rRNA gene sequences of strains WOUb02^T^ and WOUb67^T^ were found across various environments. The top detection rates are listed as follows for WOUb02^T^: human gut (75.3% of samples, with a mean relative abundance of 7.9%), insect gut (62.5%, 3.8%), rhizosphere (64.9%, 1.3%), and plant-associated datasets (59.7%, 3.9%). For WOUb67^T^ datasets from rhizosphere (74.6%, 7.5%), wastewater (67.2%, 1.0%), plant-associated (64.2%, 2.8%), and freshwater-associated (59.5%, 2.9%) were reflective of the top detection rates. Collectively, the high prevalence in plant-associated metagenomes suggests that WOUb02^T^ and WOUb67^T^ are well-adapted to persist in plant- and insect-associated ecologies.

WOUb02^T^ was isolated from a *F. septica* syconium, which represents the natural ecological niche of *C. inopinata* nematodes. These nematodes use bacteria as a food source. As WOUb02^T^ and *C. inopinata* were originally isolated from the same plant host, we recently described how *C. inopinata* responds to WOUb02^T^ in laboratory conditions [17]. We found that WOUb02^T^ doubles the population-level growth rate of *C. inopinata* in laboratory conditions (compared with the traditional food source of *E. coli*); this fitness increase is largely due to the increased developmental rate of *C. inopinata* when reared with WOUb02^T^. Moreover, we performed 16S rRNA high-throughput metabarcoding (n=70) of fig samples [18], revealing that WOUb02^T^ is widely found in *F. septica* figs [17]. While WOUb02^T^ showed no association with foundress pollinating wasp number or worm occupancy, WOUb02^T^ abundance was higher in fig interiors compared to fig exteriors [17, 18]. Thus, WOUb02^T^ is likely associated with interior syconium structures such as the reproductive organs or the fig lumen. Regardless, as *Klebsiella* spp. are frequently associated with *Caenorhabditis* nematodes [21, 23, 47, 48], and as WOUb02^T^ increases both the fitness of *C. inopinata* and *C. elegans* [17], WOUb02^T^ is likely to represent one of myriad novel bacterial species meaningfully interacting with these animals.

WOUb67^T^ was isolated from a *F. benguetensis* syconium. Although the microbial communities of this fig species have not been characterized with high-throughput metabarcoding, previous studies have revealed that *Pseudomonas* spp. are commonly found in plant-associated substrates [21, 23, 47]. Indeed, 40 *Pseudomonas* amplicon sequence variants were detected in *F. septica* figs alone [49], and six of these harbored >98.0% identity to the 16S sequence of WOUb67^T^. Thus, much like WOUb2^T^, WOUb67^T^ is likely a member of a lineage of microbes with a deep evolutionary history with plants and their associated free-living nematodes.

### Physiological analysis

Unless otherwise stated, all tests were performed in triplicate. The Gram stain reaction was performed using the Advanced Gram Stain Kit (Hardy Diagnostics, Santa Maria, CA, USA) according to the manufacturer’s instructions. The strain’s relationship to oxygen was determined by incubating cultures at 37°C for 72 hours in a fluid thioglycollate medium (Becton, Dickinson and Company, Franklin Lakes, NJ). Cells were inspected for motility and endospore production by phase contrast microscopy (CX41, Olympus, Center Valley, PA, USA) at x1,000 magnification. Cells were also examined for motility by observing growth in motility test medium as previously described [50]. The temperature range for growth was determined in tryptic soy broth (TSB) at 4°C, 20°C, 25°C, 30°C, 37°C, 45°C, 50°C, and 55°C. Salt tolerance was determined in TSB at 0.5% NaCl (w/v) and between 0.0% NaCl (w/v) and 10.0% NaCl (w/v), in increments of 1.0% (w/v); pH ranges for growth were determined between 4.0 and 10.0, in increments of 1.0 in TSB. For the pH ranges, buffers were prepared at a final concentration of 10 mM as previously described [51]. Optimal growth conditions were determined by optical density at 600nm using a BioTek 800 TS Absorbance Reader (Agilent, Santa Clara, CA, USA). An increase in optical density greater than 0.1 after five days of incubation was considered growth. The following conventional phenotypic tests: indole, citrate, Methyl Red, Voges-Proskauer, urease, and nitrate reduction were also performed for both strains as previously described [52]. Additionally, for strain WOUb67^T^, fluorescence compound production was determined using King’s A and King’s B media, as described by King *et al.* (1954) [53]. Additional biochemical characterization of isolates WOUb02^T^ and WOUb67^T^ was performed according to the manufacturer’s instructions using a Biolog GEN III MicroPlate (BIOLOG, Hayward, CA, USA) and scored by eye after 24 hours of incubation at 30°C. Detailed phenotypic characteristics of WOUb02^T^ and WOUb67^T^ are provided in their respective species descriptions. Distinguishing phenotypic characteristics between strains WOUb02^T^ and WOUb67^T^ and close relatives are listed in Tables 3 and 4.

**Table 3.**
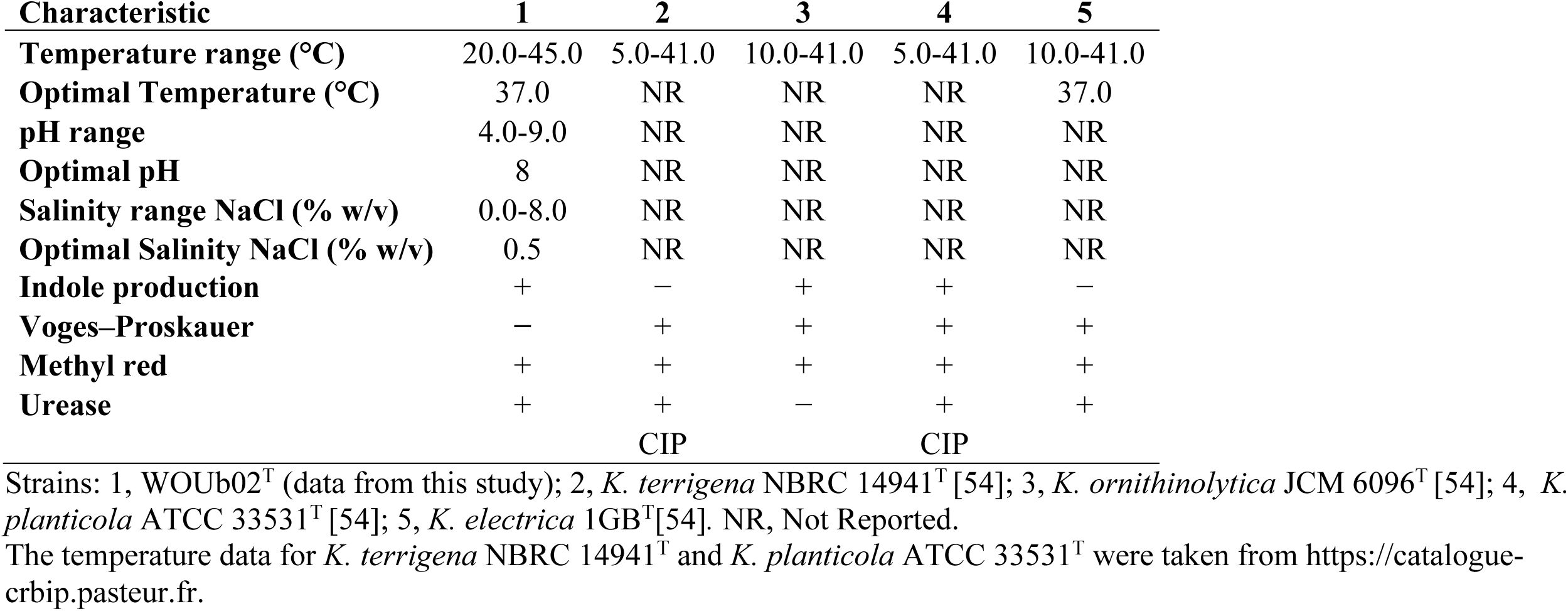
Physiological differential characteristics of WOUb02^T^ and close relatives.

| **Characteristic** | **1** | **2** | **3** | **4** | **5** |
|---|---|---|---|---|---|
| **Temperature range (°C)** | 20.0-45.0 | 5.0-41.0 | 10.0-41.0 | 5.0-41.0 | 10.0-41.0 |
| **Optimal Temperature (°C)** | 37.0 | NR | NR | NR | 37.0 |
| **pH range** | 4.0-9.0 | NR | NR | NR | NR |
| **Optimal pH** | 8 | NR | NR | NR | NR |
| **Salinity range NaCl (% w/v)** | 0.0-8.0 | NR | NR | NR | NR |
| **Optimal Salinity NaCl (% w/v)** | 0.5 | NR | NR | NR | NR |
| **Indole production** | + | − | + | + | − |
| **Voges–Proskauer** | − | + | + | + | + |
| **Methyl red** | + | + | + | + | + |
| **Urease** | + | + | − | + | + |
|  |  | CIP |  | CIP |  |

Strains: 1, WOUb02<sup>T</sup> (data from this study); 2, *K. terrigena* NBRC 14941<sup>T</sup> [54]; 3, *K. ornithinolytica* JCM 6096<sup>T</sup> [54]; 4,  *K. planticola* ATCC 33531<sup>T</sup> [54]; 5, *K. electrica* 1GB<sup>T</sup>[54]. NR, Not Reported.
The temperature data for *K. terrigena* NBRC 14941<sup>T</sup> and *K. planticola* ATCC 33531<sup>T</sup> were taken from https://catalogue-crbip.pasteur.fr.

**Table 4.** Physiological differential characteristics of WOUb67^T^ and close relatives.

| **Characteristic** | **1** | **2** | **3** | **4** | **5** |
|---|---|---|---|---|---|
| **Temperature range (°C)** | 20-37 | NR | NR | NR | NR |
| **Optimal Temperature (°C)** | 30 | NR | NR | NR | NR |
| **pH range** | 5.0-10.0 | NR | NR | NR | NR |
| **Optimal pH** | 7 | NR | NR | NR | NR |
| **Salinity range NaCl (% w/v)** | 0.0-3.0 | NR | NR | NR | NR |
| **Optimal Salinity NaCl (% w/v)** | 0.5 | NR | NR | NR | NR |
| **Utilization of:** |  |  |  |  |  |
| **Propionic acid** | - | + | + | + | + |
| **Acetic acid** | - | + | + | + | + |
| **Formic acid** | - | + | + | + | - |
| **L-Alanine** | - | + | + | + | + |
| **L-Serine** | - | + | + | + | + |
| **D-Gluconic acid** | - | + | + | + | + |

Strains: 1, WOUb67<sup>T</sup> (data from this study); 2, *P. putida* NBRC 14164<sup>T</sup> [55]; 3, *P. promysalinigenes* RW10S1<sup>T</sup> [16]; 4, *P. kermanshahensis* SWR100<sup>T</sup> [16]; 5, *P. urmiensis* SWRI10<sup>T</sup> [16]. NR, Not Reported.

The cells of strain WOUb02^T^ are Gram-stain-negative and facultatively anaerobic. Colonies were white and circular on the surface of TSA after 24 hours at 37°C. Cells were non-motile and endospores are not observed. Strain WOUb02^T^ could grow between 20°C and 45°C with optimum growth at 37°C, growth was observed between 0.0% and 8.0% (w/v) NaCl concentrations, with optimal growth at 0.5% (w/v). Isolate WOUb02^T^ grew at pH values between pH 4.0 and 9.0, with an optimum pH of 8.0. It was positive for indole, citrate, Methyl Red, urease, and nitrate reduction; Voges-Proskauer produced a negative result. Strain WOUb02^T^ exhibited a broad substrate utilization profile, including mono-, di-, and oligosaccharides (e.g., D-glucose, D-maltose, D-cellobiose, raffinose, and stachyose), amino sugars (e.g., N-acetyl-D-glucosamine and N-acetylneuraminic acid), sugar alcohols (e.g., D-mannitol and myo-inositol), and organic acids (e.g., citric acid and D-malic acid). The strain also utilized pectin and related uronic acids, indicating the capacity for complex polysaccharide degradation. Additionally, WOUb02^T^ displayed tolerance to multiple antibiotics and inhibitory compounds, including rifamycin SV, minocycline, and vancomycin. Strain WOUb02ᵀ can be distinguished from *K. terrigena* by positive indole production (Table 3). Notably, WOUb02^T^ was Voges–Proskauer negative (Table 3).

The cells of strain WOUb67^T^ are Gram-stain-negative and was obligately aerobic. Colonies were mucoid on TSA after 24 hours at 30°C. Cells were motile and endospores were not observed. Strain WOUb67^T^ could grow between 20°C and 37°C, with optimum growth at 30°C. It could grow at NaCl concentrations between 0.0% and 3.0% (w/v), with optimal growth at 0.5% (w/v). Growth was observed at pH values between 5.0 and 10.0, with an optimum pH of 7.0. Positive reactions were obtained for citrate, urease, and nitrate reduction; indole, Methyl Red, and Voges-Proskauer produced negative results.

Strain WOUb67ᵀ exhibited a more restricted substrate utilization profile, including select monosaccharides (e.g., α-D-glucose, D-fructose, and D-/L-fucose), amino acids (e.g., L-arginine, L-aspartic acid, L-glutamic acid, L-histidine, and L-pyroglutamic acid), and organic acids (e.g., citric acid and L-malic acid). The strain also utilized uronic acid derivatives, including D-galacturonic acid, D-glucuronic acid, and glucuronamide. Additionally, WOUb67^T^ strain demonstrated tolerance to multiple antibiotics and inhibitory compounds, including rifamycin SV, vancomycin, nalidixic acid, and aztreonam, as well as chemical stressors such as lithium chloride and potassium tellurite. Notably, WOUb67^T^ differs from its closest neighbors in that it is unable to utilize propionic acid, acetic acid, formic acid, L-alanine, L-serine, and D-gluconic acid as sole carbon sources (Table 4).

### Taxonomic Conclusions

Based on polyphasic taxonomic criteria, WOUb02^T^ and WOUb67^T^ represent novel species within *Klebsiella* and *Pseudomonas,* respectively. Therefore, we assign strains WOUb02^T^ and WOUb67^T^ as novel species in the genera *Klebsiella* and *Pseudomonas* in this study, for which the names *Klebsiella taiwanensis* (=NRRL-B 65719^T^ =CCUG 77855^T^) sp. nov. and *Pseudomonas ficus* (=NRRL-B 65720^T^ =CCUG 77856^T^sp. nov., are proposed.

## PROTOLOGUE

### DESCRIPTION OF KLEBSIELLA TAIWANENSIS SP. NOV

*Klebsiella taiwanensis* (tai.wa.nen’sis. N.L. fem. adj. *taiwanensis,* pertaining to Taiwan, where the type strain was isolated). The cells are Gram-stain-negative rods, facultatively anaerobic, non-motile, sporulation is not observed. Colonies on tryptic soy agar medium are white, dome-shaped, and sticky on the agar surface after 24 hours at 37°C. Optimal growth is observed at 37°C, with 0.5% (w/v) NaCl, at pH 8.0. Positive for growth between 20°C and 45°C. Positive for growth between 0.0% and 8.0% (w/v) NaCl and between pH 4.0 and pH 9.0. Positive results are obtained for the indole, citrate, Methyl Red, urease, and nitrate reductase tests; Voges-Proskauer test is negative. Cells utilize Dextrin, D-Maltose, D-Trehalose, D-Cellobiose, Gentiobiose, D-Turanose, Stachyose, D-Raffinose, α-D-Lactose, D-Melibiose, β-Methyl-D-Glucoside, N-Acetyl-D-Glucosamine, N-Acetyl-β-D-Mannosamine, N-Acetyl-D-Galactosamine, N-Acetyl Neuraminic Acid, α-D-Glucose, D-Mannose, D-Fructose, D-Galactose, D-Fucose, L-Fucose, L-Rhamnose, Inosine, 1% Sodium Lactate, D-Serine, D-Sorbitol, D-Mannitol, D-Arabitol, myo-Inositol, Glycerol, D-Glucose-6-PO₄, D-Fructose-6-PO₄, Glycyl-L-Proline, L-Alanine, L-Arginine, L-Aspartic Acid, L-Glutamic Acid, L-Histidine, L-Serine, Pectin, D-Galacturonic Acid, L-Galactonic Acid Lactone, D-Gluconic Acid, D-Glucuronic Acid, Glucuronamide, Mucic Acid, D-Saccharic Acid, p-Hydroxy-Phenylacetic Acid, Methyl Pyruvate, L-Lactic Acid, Citric Acid, D-Malic Acid, L-Malic Acid, Bromo-Succinic Acid, Acetic Acid, Formic Acid, and Sodium Butyrate. Cells grow in the presence of Fusidic Acid, Rifamycin SV, Minocycline, Troleandomycin, Lincomycin, Vancomycin, Nalidixic Acid, Aztreonam, Lithium Chloride, Guanidine HCl, Niaproof 4, Tetrazolium Violet, and Tetrazolium Blue.

The type strain is WOUb02^T^ (=NRRL-B 65719^T^ =CCUG 77855^T^), which was isolated from the homogenate of fresh fig, *Ficus septica*, sampled in Taipei, Taiwan (25.0314672 N, 121.5749889 E). The G+C content is 57.0%. The 16S rRNA gene and genome sequences of WOUb02^T^ have been deposited in GenBank under the accession numbers PP890003.1 and GCA_055451655.1, respectively.

### DESCRIPTION OF PSEUDOMONAS FICUS SP. NOV

*Pseudomonas ficus* (fi’cus. L. gen. n. *ficus,* of the fig tree, of the genus *Ficus*., referring to the source of isolation for the type strain). The cells are Gram-stain-negative rods, strictly aerobic, and motile, sporulation is not observed. Colonies on tryptic soy agar are mucoid on the agar surface after 24 hours at 30°C. Optimal growth is observed at 30°C, with 0.5% (w/v) NaCl, at pH 7.0. Positive for growth between 20°C and 37°C. Positive for growth between 0.0% and 3.0% (w/v) NaCl and between pH 5.0 and pH 10.0. Positive results are obtained for the citrate, urease, and nitrate reduction tests; Indole, Methyl Red, and Voges-Proskauer tests is negative. Cells utilize α-D-Glucose, D-Fructose, D-Fucose, L-Fucose, 1% Sodium Lactate, D-Serine, D-Aspartic Acid, L-Arginine, L-Aspartic Acid, L-Glutamic Acid, L-Histidine, L-Pyroglutamic Acid, D-Galacturonic Acid, L-Galactonic Acid Lactone, D-Glucuronic Acid, Glucuronamide, Mucic Acid, Quinic Acid, p-Hydroxy-Phenylacetic Acid, Citric Acid, L-Malic Acid, and Sodium Butyrate. Cells can grow in Fusidic Acid, Troleandomycin, Rifamycin SV, Lincomycin, Guanidine HCl, Niaproof 4, Vancomycin, Tetrazolium Violet, Tetrazolium Blue, Nalidixic Acid, Lithium Chloride, Potassium Tellurite, Aztreonam, and Sodium Bromate.

The type strain is WOUb67^T^ (=NRRL-B 65720^T^=CCUG 77856^T^), which was isolated from the homogenate of fresh fig, *Ficus benguetensis*, sampled in Taipei, Taiwan (25.002000 N 121.564000 E). The G+C content is 62.5%. The 16S rRNA gene and genome sequences of WOUb67^T^ have been deposited in GenBank under the accession numbers PP902522.1 and GCA_050058005.1, respectively.

## Abbreviations

NRRL: Agricultural Research Service Culture Collection
CCUG: Culture Collection of the ANI, Average Nucleotide Identity
dDDH: Digital DNA-DNA Hybridization

## Data Accessibility

The 16S rRNA gene sequences of strains WOUb02^T^ and WOUb67^T^ were deposited in GenBank under accession numbers PP890003.1 and PP902522.1, respectively. Whole-genome sequences of strains WOUb02^T^ and WOUb67^T^ were deposited in GenBank under accession numbers GCA_055451655.1 and GCA_050058005.1, respectively. The type strain WOUb02^T^ has been deposited in the Agricultural Research Service Culture Collection (accession number: NRRL-B 65719 ^T^) and the Culture Collection University of Gothenburg (accession number: CCUG 77855^T^). The type strain WOUb67^T^ has been deposited in the Agricultural Research Service Culture Collection (accession number: NRRL-B 65720^T^) and the Culture Collection University of Gothenburg (accession number: CCUG 77865^T^).

## Ethical Statement

This article does not contain any studies with humans performed by any of the authors.

## Acknowledgments

The authors thank Professor Aharon Oren (Hebrew University of Jerusalem, Israel) for helping check the etymology of the proposed name.

---

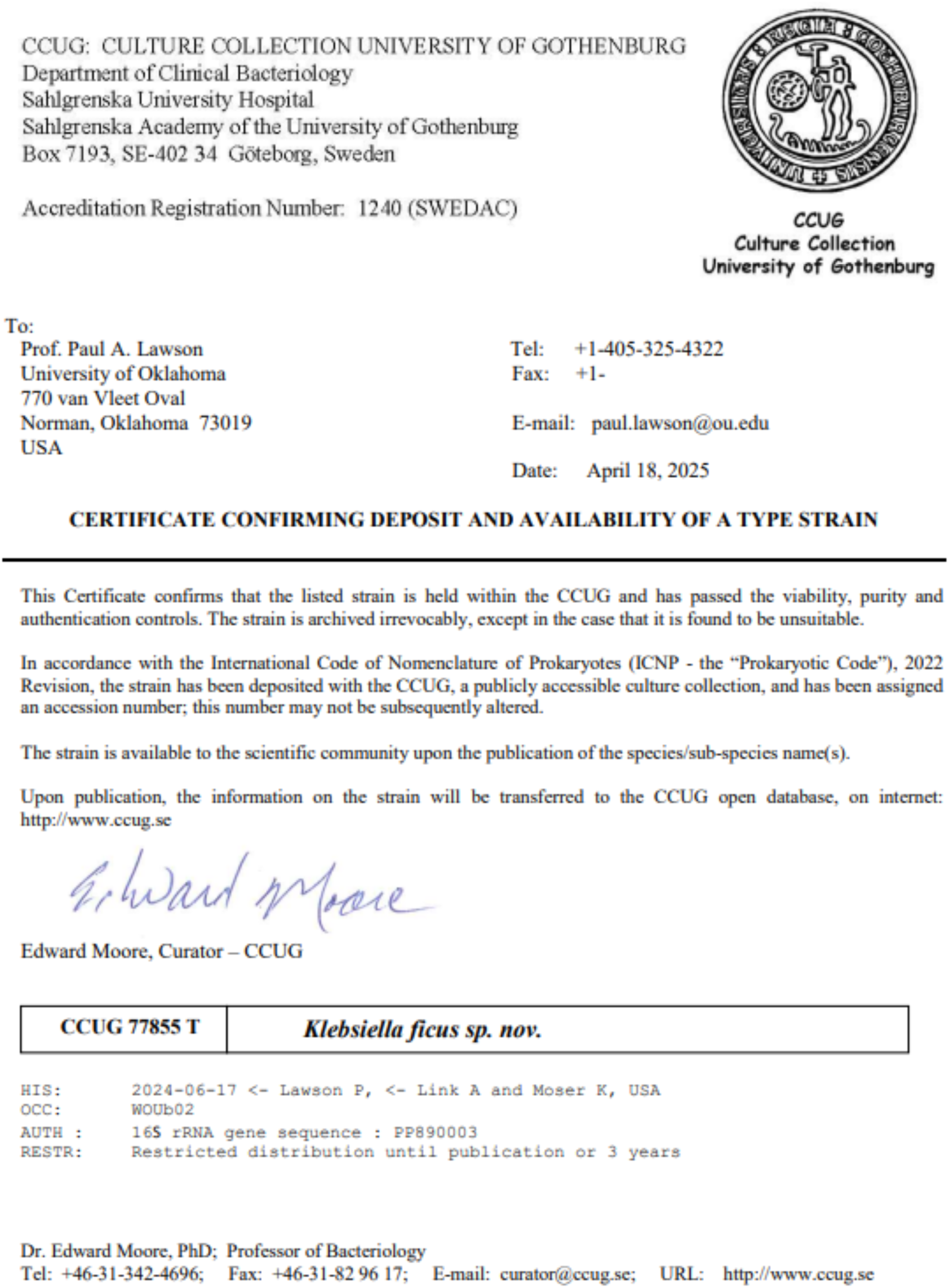

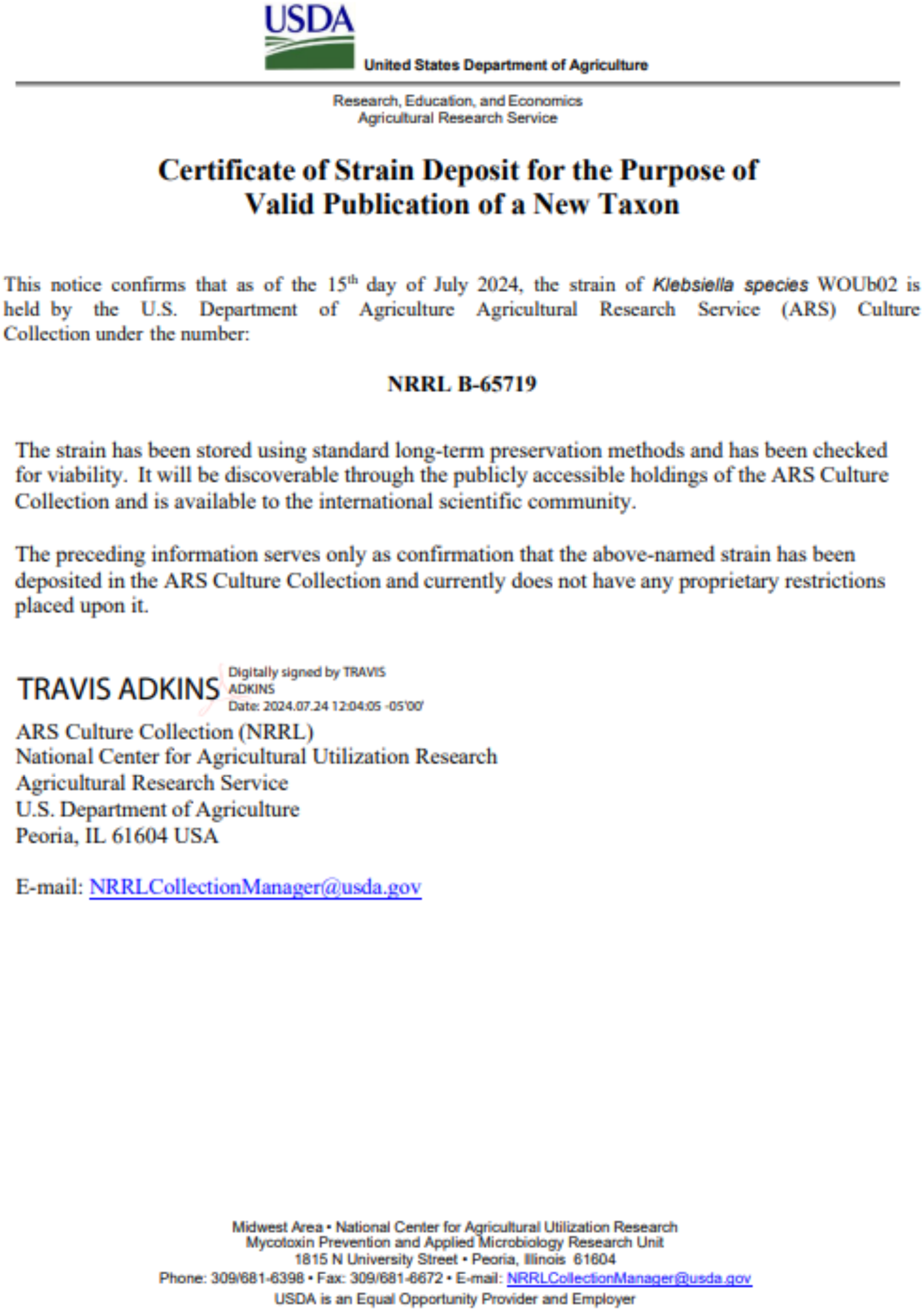

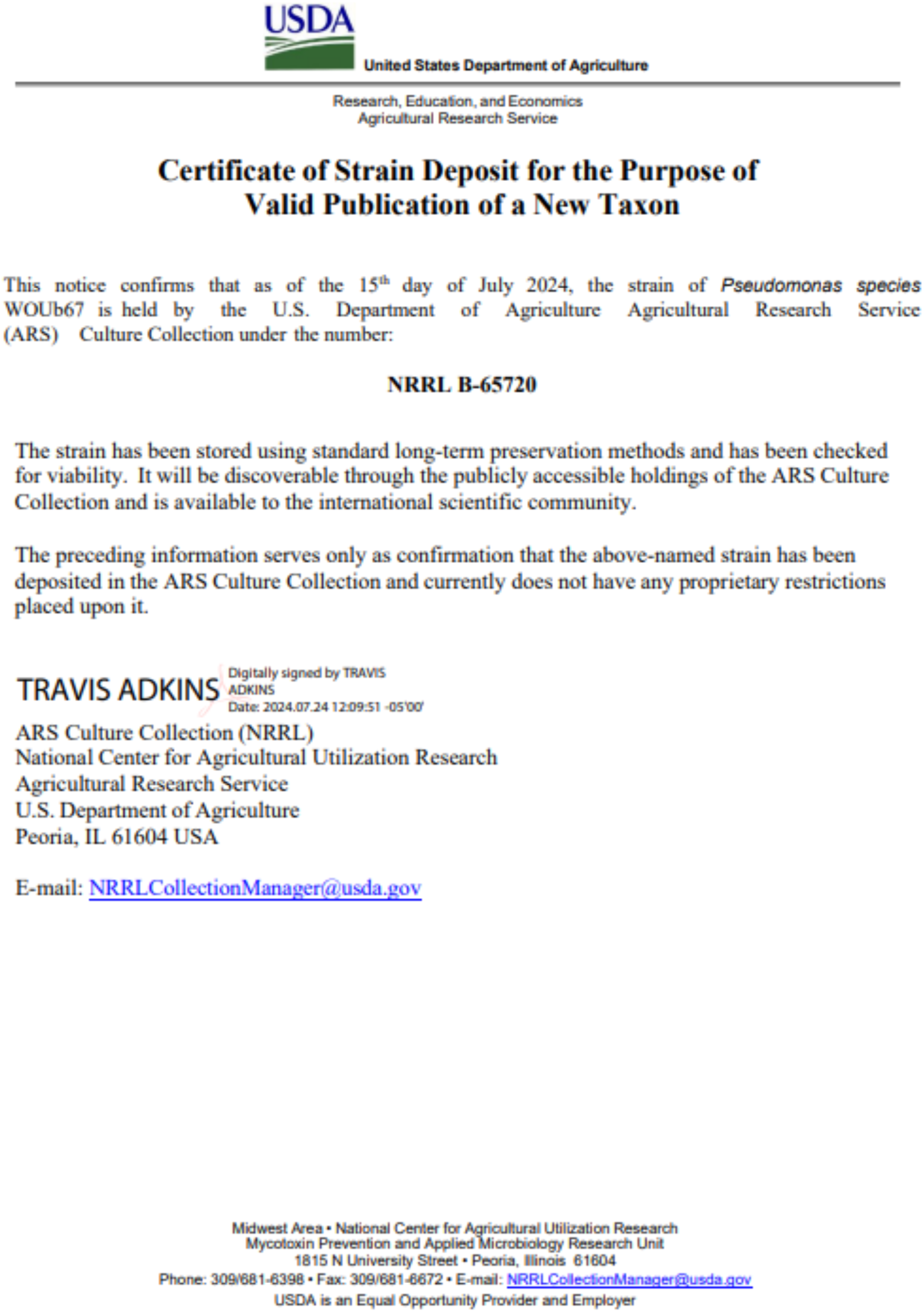

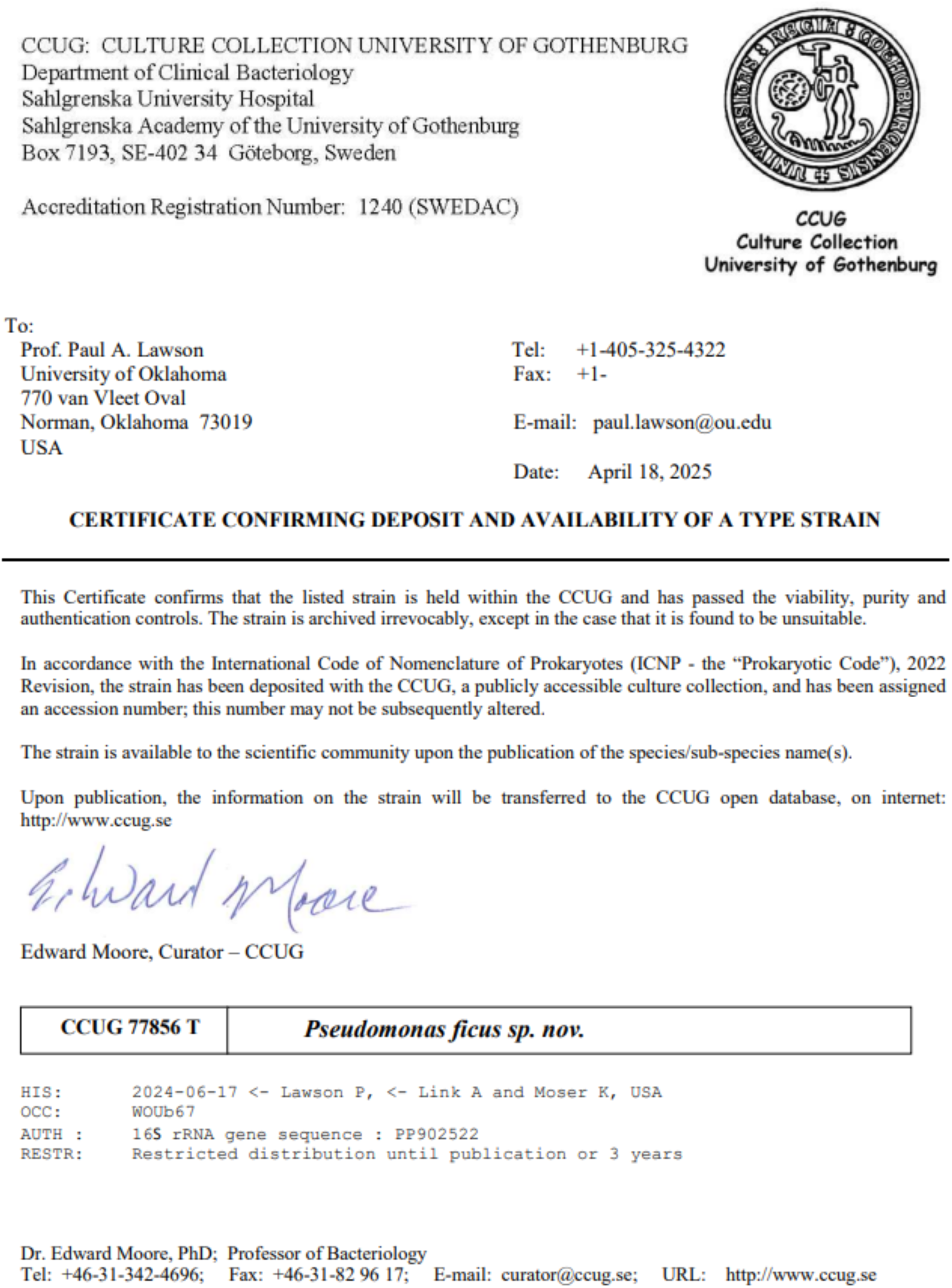

## References

1. Brenner DJ, Farmer JJ. Enterobacteriaceae. In: Whitman WB (editor). Bergey’s Manual of Systematics of Archaea and Bacteria. Wiley. pp. 1–24.

2. Dong N, Yang X, Chan EW-C, Zhang R, Chen S. Klebsiella species: Taxonomy, hypervirulence and multidrug resistance. eBioMedicine 2022;79:103998.

3. Podschun R, Ullmann U. *Klebsiella* spp. as Nosocomial Pathogens: Epidemiology, Taxonomy, Typing Methods, and Pathogenicity Factors. Clin Microbiol Rev 1998;11:589–603.

4. Li B, Zhao Y, Liu C, Chen Z, Zhou D. Molecular Pathogenesis of *Klebsiella Pneumoniae*. Future Microbiol 2014;9:1071–1081.

5. Hu Y, Anes J, Devineau S, Fanning S. *Klebsiella pneumoniae* : Prevalence, Reservoirs, Antimicrobial Resistance, Pathogenicity, and Infection: A Hitherto Unrecognized Zoonotic Bacterium. Foodborne Pathog Dis 2021;18:63–84.

6. Brisse S, Passet V, Grimont PAD. Description of *Klebsiella quasipneumoniae* sp. nov., isolated from human infections, with two subspecies, *Klebsiella quasipneumoniae* subsp. *quasipneumoniae* subsp. nov. and *Klebsiella quasipneumoniae* subsp. *similipneumoniae* subsp. nov., and demonstration that *Klebsiella singaporensis* is a junior heterotypic synonym of *Klebsiella variicola*. Int J Syst Evol Microbiol 2014;64:3146–3152.

7. Rodríguez-Medina N, Barrios-Camacho H, Duran-Bedolla J, Garza-Ramos U. *Klebsiella variicola* : an emerging pathogen in humans. Emerg Microbes Infect 2019;8:973–988.

8. Hu Y, Wei L, Feng Y, Xie Y, Zong Z. Klebsiella huaxiensis sp. nov., recovered from human urine. Int J Syst Evol Microbiol 2019;69:333–336.

9. Izard D, Ferragut C, Gavini F, Kersters K, De Ley J, et al. *Klebsiella terrigena*, a New Species from Soil and Water. Int J Syst Bacteriol 1981;31:116–127.

10. Ferragut C, Izard D, Gavini F, Kersters K, De Ley J, et al. *Klebsiella trevisanii*: a New Species from Water and Soil. Int J Syst Bacteriol 1983;33:133–142.

11. Gujarati S, Chaudhari D, Hagir A, Khairnar M, Shouche Y, et al. *Klebsiella indica* sp. nov., isolated from the surface of a tomato. Int J Syst Evol Microbiol 2020;70:3278–3286.

12. Rosenblueth M, Martínez L, Silva J, Martínez-Romero E. *Klebsiella variicola*, A Novel Species with Clinical and Plant-Associated Isolates. Syst Appl Microbiol 2004;27:27–35.

13. Palleroni NJ. Pseudomonas. In: Whitman WB (editor). Bergey’s Manual of Systematics of Archaea and Bacteria. Wiley. pp. 1–1.

14. De Bentzmann S, Plésiat P. The *Pseudomonas aeruginosa* opportunistic pathogen and human infections. Environ Microbiol 2011;13:1655–1665.

15. Azam MW, Khan AU. Updates on the pathogenicity status of *Pseudomonas aeruginosa*. Drug Discov Today 2019;24:350–359.

16. Girard L, Lood C, Höfte M, Vandamme P, Rokni-Zadeh H, et al. The Ever-Expanding *Pseudomonas* Genus: Description of 43 New Species and Partition of the *Pseudomonas putida* Group. Microorganisms 2021;9:1766.

17. Link AC, Moser KA, Wang J, Woodruff GC. A novel, fig-associated microbe promotes reproductive success via variable life history mechanisms in Caenorhabditis elegans and *C. inopinata*. bioRxiv 2025. doi:10.64898/2025.12.16.694684.

18. Woodruff GC, Moser KA, Wang J. The bacteria of a fig community. Microbiol Spectr 2026;14:e03013–24.

19. Kiontke KC, Félix M-A, Ailion M, Rockman MV, Braendle C, et al. A phylogeny and molecular barcodes for *Caenorhabditis*, with numerous new species from rotting fruits. BMC Evol Biol 2011;11:339.

20. Félix M-A, Duveau F. Population dynamics and habitat sharing of natural populations of *Caenorhabditis elegans* and *C. briggsae*. BMC Biol 2012;10:59.

21. Dirksen P, Marsh SA, Braker I, Heitland N, Wagner S, et al. The native microbiome of the nematode *Caenorhabditis elegans*: gateway to a new host-microbiome model. BMC Biol 2016;14:38.

22. Crombie TA, Zdraljevic S, Cook DE, Tanny RE, Brady SC, et al. Deep sampling of Hawaiian *Caenorhabditis elegans* reveals high genetic diversity and admixture with global populations. eLife 2019;8:e50465.

23. Samuel BS, Rowedder H, Braendle C, Félix M-A, Ruvkun G. *Caenorhabditis elegans* responses to bacteria from its natural habitats. Proc Natl Acad Sci U S A 2016;113–E3949. doi:10.1073/pnas.1607183113.

24. Kanzaki N, Tsai IJ, Tanaka R, Hunt VL, Liu D, et al. Biology and genome of a newly discovered sibling species of *Caenorhabditis elegans*. Nat Commun 2018;9:3216.

25. Woodruff GC, Phillips PC. Field studies reveal a close relative of *C. elegans* thrives in the fresh figs of *Ficus septica* and disperses on its Ceratosolen pollinating wasps. BMC Ecol 2018;18:26.

26. Brenner S. The Genetics of *Caenorhabditis elegans*. Genetics 1974;77:71–94.

27. Yoon S-H, Ha S-M, Kwon S, Lim J, Kim Y, et al. Introducing EzBioCloud: a taxonomically united database of 16S rRNA gene sequences and whole-genome assemblies. Int J Syst Evol Microbiol 2017;67:1613–1617.

28. Edgar RC. MUSCLE: multiple sequence alignment with high accuracy and high throughput. Nucleic Acids Res 2004;32:1792–1797.

29. Felsenstein J. Evolutionary trees from DNA sequences: A maximum likelihood approach. J Mol Evol 1981;17:368–376.

30. Tamura K, Nei M. Estimation of the number of nucleotide substitutions in the control region of mitochondrial DNA in humans and chimpanzees. Mol Biol Evol 1993; 10:512–526

31. Kumar S, Stecher G, Suleski M, Sanderford M, Sharma S, et al. MEGA12: Molecular Evolutionary Genetic Analysis Version 12 for Adaptive and Green Computing. Mol Biol Evol 2024;41:msae263.

32. Kim M, Oh H-S, Park S-C, Chun J. Towards a taxonomic coherence between average nucleotide identity and 16S rRNA gene sequence similarity for species demarcation of prokaryotes. Int J Syst Evol Microbiol 2014;64:346–351.

33. Mulet M, Lalucat J, García-Valdés E. DNA sequence-based analysis of the *Pseudomonas* species. Environ Microbiol 2010;12:1513–1530.

34. Miller SL, Youssef NH, Elshahed MS. Genome-based reclassification of *Parabacteroides chartae* as a later heterotypic synonym of *Macellibacteroides fermentans*. Int J Syst Evol Microbiol 2025; 75

35. Parks DH, Imelfort M, Skennerton CT, Hugenholtz P, Tyson GW. CheckM: assessing the quality of microbial genomes recovered from isolates, single cells, and metagenomes. Genome Res 2015;25:1043–1055.

36. Tatusova T, DiCuccio M, Badretdin A, Chetvernin V, Nawrocki EP, et al. NCBI prokaryotic genome annotation pipeline. Nucleic Acids Res 2016;44:6614–6624.

37. Riesco R, Trujillo ME. Update on the proposed minimal standards for the use of genome data for the taxonomy of prokaryotes. Int J Syst Evol Microbiol 2024; 74:006300

38. Yoon S-H, Ha S, Lim J, Kwon S, Chun J. A large-scale evaluation of algorithms to calculate average nucleotide identity. Antonie Van Leeuwenhoek 2017;110:1281–1286.

39. Meier-Kolthoff JP, Auch AF, Klenk H-P, Göker M. Genome sequence-based species delimitation with confidence intervals and improved distance functions. BMC Bioinformatics 2013;14:60.

40. Meier-Kolthoff JP, Carbasse JS, Peinado-Olarte RL, Göker M. TYGS and LPSN: a database tandem for fast and reliable genome-based classification and nomenclature of prokaryotes. Nucleic Acids Res 2022;50:D801–D807.

41. Letunic I, Bork P. Interactive Tree of Life (iTOL) v6: recent updates to the phylogenetic tree display and annotation tool. Nucleic Acids Res 2024;52:W78–W82.

42. Richter M, Rosselló-Móra R. Shifting the genomic gold standard for the prokaryotic species definition. Proc Natl Acad Sci 2009;106:19126–19131.

43. Yin Y, Mao X, Yang J, Chen X, Mao F, et al. dbCAN: a web resource for automated carbohydrate-active enzyme annotation. Nucleic Acids Res 2012;40:W445–W451.

44. Zheng J, Ge Ǫ, Yan Y, Zhang X, Huang L, et al. dbCAN3: automated carbohydrate-active enzyme and substrate annotation. Nucleic Acids Res 2023;51:W115–W121.

45. Hitch TCA, Riedel T, Oren A, Overmann J, Lawley TD, et al. Automated analysis of genomic sequences facilitates high-throughput and comprehensive description of bacteria. ISME Commun 2021;1:16.

46. Lagkouvardos I, Joseph D, Kapfhammer M, Giritli S, Horn M, et al. IMNGS: A comprehensive open resource of processed 16S rRNA microbial profiles for ecology and diversity studies. Sci Rep 2016;6:33721.

47. Zhang F, Weckhorst JL, Assié A, Hosea C, Ayoub CA, et al. Natural genetic variation drives microbiome selection in the Caenorhabditis elegans gut. Curr Biol 2021;31:2603–2618.e9.

48. Lee S, Kim Y, Choi J. Effect of soil microbial feeding on gut microbiome and cadmium toxicity in Caenorhabditis elegans. Ecotoxicol Environ Saf 2020;187:109777.

49. Woodruff GC, Moser KA, Wang J. The bacteria of a fig community. Microbiol Spectr 2026;e03013–24.

50. Jain A, Jain R, Jain S. Motility Testing – Hanging Drop Method and Stab. In: Jain A, Jain R, Jain S (editors). Basic Techniques in Biochemistry, Microbiology and Molecular Biology: Principles and Techniques. New York, NY: Springer US. pp. 121–122.

51. Miller S, Hendry M, King J, Sankaranarayanan K, Lawson PA. *Clostridium tanneri* sp. nov., isolated from the faecal material of an alpaca. Int J Syst Evol Microbiol 2024; 74:006372

52. Lányi B. 1 Classical and Rapid Identification Methods for Medically Important Bacteria. In: Methods in Microbiology. Elsevier. pp. 1–67.

53. King EO, Ward MK, Raney DE. Two simple media for the demonstration of pyocyanin and fluorescin. J Lab Clin Med 1954;44:301–307.

54. Rahi P, Rodrigues C, Brisse S. Klebsiella. In: Trujillo ME, Dedysh S, DeVos P, Hedlund B, Kämpfer P, et al. (editors). Bergey’s Manual of Systematics of Archaea and Bacteria. Wiley. pp. 1–23.

55. Mulet M, Gomila M, Lemaitre B, Lalucat J, García-Valdés E. Taxonomic characterisation of *Pseudomonas* strain L48 and formal proposal of *Pseudomonas entomophila* sp. nov. Syst Appl Microbiol 2012;35:145–149.

